# Generation of eco-friendly channel catfish, *Ictalurus punctatus*, harboring alligator cathelicidin gene with robust disease resistance by harnessing different CRISPR/Cas9-mediated systems

**DOI:** 10.1101/2023.01.05.522889

**Authors:** Jinhai Wang, Baofeng Su, De Xing, Timothy J. Bruce, Shangjia Li, Logan Bern, Mei Shang, Andrew Johnson, Rhoda Mae C. Simora, Michael Coogan, Darshika U. Hettiarachchi, Wenwen Wang, Tasnuba Hasin, Jacob Al-Armanazi, Cuiyu Lu, Rex A. Dunham

## Abstract

The CRISPR/Cas9 platform holds promise for modifying fish traits of interest as a precise and versatile tool for genome manipulation. To reduce introgression of transgene and control reproduction, catfish species have been studied for upscaled disease resistance and intervening of reproduction to lower the potential environmental risks of introgression of escapees’ as transgenic animals. Taking advantage of the CRISPR/Cas9-mediated system, we succeeded in integrating the cathelicidin gene from an alligator (*Alligator sinensis*; *As-Cath*) into the target luteinizing hormone (*LH*) locus of channel catfish (*Ictalurus punctatus*) using two delivery systems assisted by double-stranded DNA (dsDNA) and single-stranded oligodeoxynucleotides (ssODNs), respectively. In this study, high knock-in (KI) efficiency (22.38%, 64/286) but low on-target was achieved using the ssODN strategy, whereas adopting a dsDNA as the donor template led to an efficient on-target KI (10.80%, 23/213). On-target KI of *As-Cath* was instrumental in establishing the *LH* knockout (LH^−^_As-Cath^+^) catfish line, which displayed heightened disease resistance and reduced fecundity compared to the wild-type sibling fish. Furthermore, implanting with HCG and LHRHa can restore the fecundity, spawnability and hatchability of the new transgenic fish line. Overall, we replaced the *LH* gene with an alligator cathelicidin transgene and then administered hormone therapy to gain complete reproductive control of disease-resistant transgenic catfish in an environmentally sound manner. This strategy not only effectively improves the consumer-valued traits, but also guards against genetic contamination. This is a breakthrough in aquaculture genetics to confine fish reproduction and prevent the establishment of transgenic or domestic genotypes in the natural environment.

## 1. Introduction

Innovative biotechnologies continuously develop as science advances, benefiting food production, quality as well as animal and human welfare. Since its inception, CRISPR/Cas9 (clustered regularly interspaced short palindromic repeats/CRISPR-associated protein 9) has served as a prototype in genome engineering, paving the way for new possibilities in transgenesis and breeding. Two mechanisms are involved for DNA repair when double strand breaks are induced by the CRISPR/Cas9 complex: non-homologous end joining (NHEJ) and homology-directed repair (HDR) [1]. Both mechanism-mediated strategies have been employed in aquaculture to improve the consumer-valued qualities targeted within genetic breeding programs. These harness the NHEJ repair pathway to knock out (KO)/disrupt functional genes or knock in (KI) exogenous genes of interest via HDR at the expected locus to improve the target traits.

Numerous CRISPR/Cas9 systems have emerged recently to improve target-editing efficiency for KI via the HDR pathway. Success has been observed in model animals have been shown successes using ssODN-mediated KIs for the targeted insertions of small DNA fragments since single-stranded oligodeoxynucleotides (ssODNs) act as templates for repairing DNA damage [2-4]. Yoshimi et al. [5] have optimized the ssODN-mediated approach to knock-in larger sequences by the combination of CRISPR/Cas9 system with two 80-bp ssODNs in length. In contrast to conventional plasmid donors, the donor vector used in this system does not require homologous arms (HAs), enabling the insertion of a large vector (CAG-GFP, 4.8 kb) into the designated site (*rRosa26*) with a ∼10% integration rate in rat zygotes [5]. Later, using the CRISPR/Cas9-ssODNs mediated KI system, a 10.96% KI efficiency in sheep zygotes was determined [6]. Boel et al. [7] first applied this optimized system to a fish model, zebrafish (*Danio rerio*), and sequencing results revealed that erroneous repair was more likely to occur when ssODNs were used as repair templates. Alternatively, the modified donor plasmid containing two HAs flanked by two single guide (sgRNA)-targeted sequences (double-cut donors) typically results in a site-specific KI with a high integration rate [8,9], and this HA-medicated KI has been adapted to zebrafish and medaka (*Oryzias latipes*) [9,10]. Theoretically, if we directly offer a linear double-stranded DNA (dsDNA) flanked by two HAs derived from 5’- and 3’-ends of the targeted site and ignore the difference in stability between circular DNA and dsDNA donors, the KI efficiency will increase by convention. In addition to the type of donors, a proper concentration of each component of the CRISPR/Cas9 system has a great positive impact on KI by reducing off-target events and embryo lethality. In this regard, we anticipate achieving extremely efficient KI if a reliable delivery system and an optimized dosage of components are chosen in a non-model fish species.

Currently, transgenesis and CRISPR/Cas9-mediated genome editing have revolutionized traditional theories to accelerate the pace of aquaculture breeding programs, and delivered edible commercial products, such as the genetically modified AquAdvantage salmon [11,12], gene-edited tiger puffer fish and red sea bream (https://doi.org/10.1038/s41587-021-01197-8, 2022). Although the NHEJ strategy predominates in altering the consumer-focused traits of fish species, including growth, coloration, and reproduction, the HDR-mediated KI is an effective way to improve the omega-3 fatty acid content and disease resistance [13-15]. In comparison to the non-insertion of KO mutations, the integration of foreign genes by harnessing the HDR pathway usually raises concerns about low KI efficiency and introgression, which directly impact the advocacy of this method and the consumer acceptance of gene-inserted fish [16]. As a result, it is imperative to devise a strategy for both improving the desired traits and preventing introgression to alleviate public concerns about gene-inserted animals. Fortunately, numerous genome-editing-based studies have demonstrated that it is possible to render fish reproductively sterile by altering/disrupting key genes involved in reproduction via the NHEJ repair pathway. Thus, potentially reducing negative environmental effects associated with genetically modified fish [17-19]. Luteinizing hormone (*LH*) gene regulates gametogenesis and gestation through binding the receptor [20,21]. LH-deficient female zebrafish are infertile, whereas the mutant males are fertile, indicating that the *LH* gene facilitates fish oocyte maturation and triggers ovulation [22]. In addition, interruption of the *LH* gene in channel catfish and white-edged rockfish (*Sebastes taczanowskii*) can result in the production of sterile lines [21,23].

Large-scale disease outbreaks are inevitable, and methods of disease control need to be improved. Antimicrobial peptides (AMPs) are polypeptides that serve as substitutes for antibiotics in a variety of species’ initial line of defense (innate immunity) against microbial invasions without developing considerable antibiotic resistance [24,25]. AMPs and antimicrobial peptide genes (AMGs) including cecropin, hepcidin, piscidin, epinecidin-1, lysozyme, and lactoferrin have been used for decades to improve disease resistance in a variety of aquatic animals, as feed supplements or transgenes [13,26]. Cathelicidins are a particularly important AMP family, sharing the common cathelin-like domain [27] and exhibiting broad-spectrum antimicrobial and immune-modulating activities [28]. Recent investigations have shown that alligator-derived cathelicidin inhibits fish pathogens both *in vivo* and *in vitro* [29-31]. Therefore, integrating the AMG into the genomic DNA has broad prospects for establishing novel disease-resistant fish lines.

Fish transgenic for AMGs could provide a significant option to address disease problems, however, and additional goal would be to prevent the possibility of breeding of escapees with wild populations. Hypothetically, a reproductive gene such as *LH*, responsible for gametogenesis and gestation could be knocked out at the DNA level with the replacement of a cathelicidin gene, leading to sterile fish with heightened disease resistance. Genome-edited sterilized fish from this approach would have fertility temporarily restored with hormone therapy used for artificial spawning of fish, and it is achievable to produce environmentally-compatible and disease-resistant fish lines. In this study, two CRISPR/Cas9 delivery systems: HA- and ssODN-mediated KI were employed to insert the *As-Cath* gene at the channel catfish (*Ictalurus punctatus*) *LH* locus to develop a reversibly sterile and disease-resistant line. We compared the KI efficiency, hatchability and fry survival from various systems, and then restored the fertility of As-Cath-integrated sterile of P_0_ founders through hormone therapy. In addition, the bacterial resistance of P_0_ and F_1_ individuals from the new fish line was further evaluated.

## 2. Materials and methods

### 2.1. Ethical approval

The care and use of animals followed the applicable guidelines from expert training courses. Experimental protocols in the current study were approved by the Auburn University Institutional Animal Care and Use Committee (AU-IACUC). All fish studies were conducted in compliance with the procedures and standards established by the Association for Assessment and Accreditation of Laboratory Animal Care (AAALAC).

### 2.2. Target locus for gene insertion

As the target integration site, we selected the *LH* gene, which is widely expressed in the theca cells of the ovary and aids in egg maturation and ovulation during gonadal development [22]. Based on the published genome of channel catfish [32], the chosen *LH* site for sgRNA targeting was located in the middle of exon 2 (Fig. 1(A-B)). The inserted segment was derived from the coding sequence (CDS) of the cathelicidin gene of *Alligator sinensis* (*As-Cath*, GeneBank accession number XM_006037211.3) [29].

**Fig. 1.**
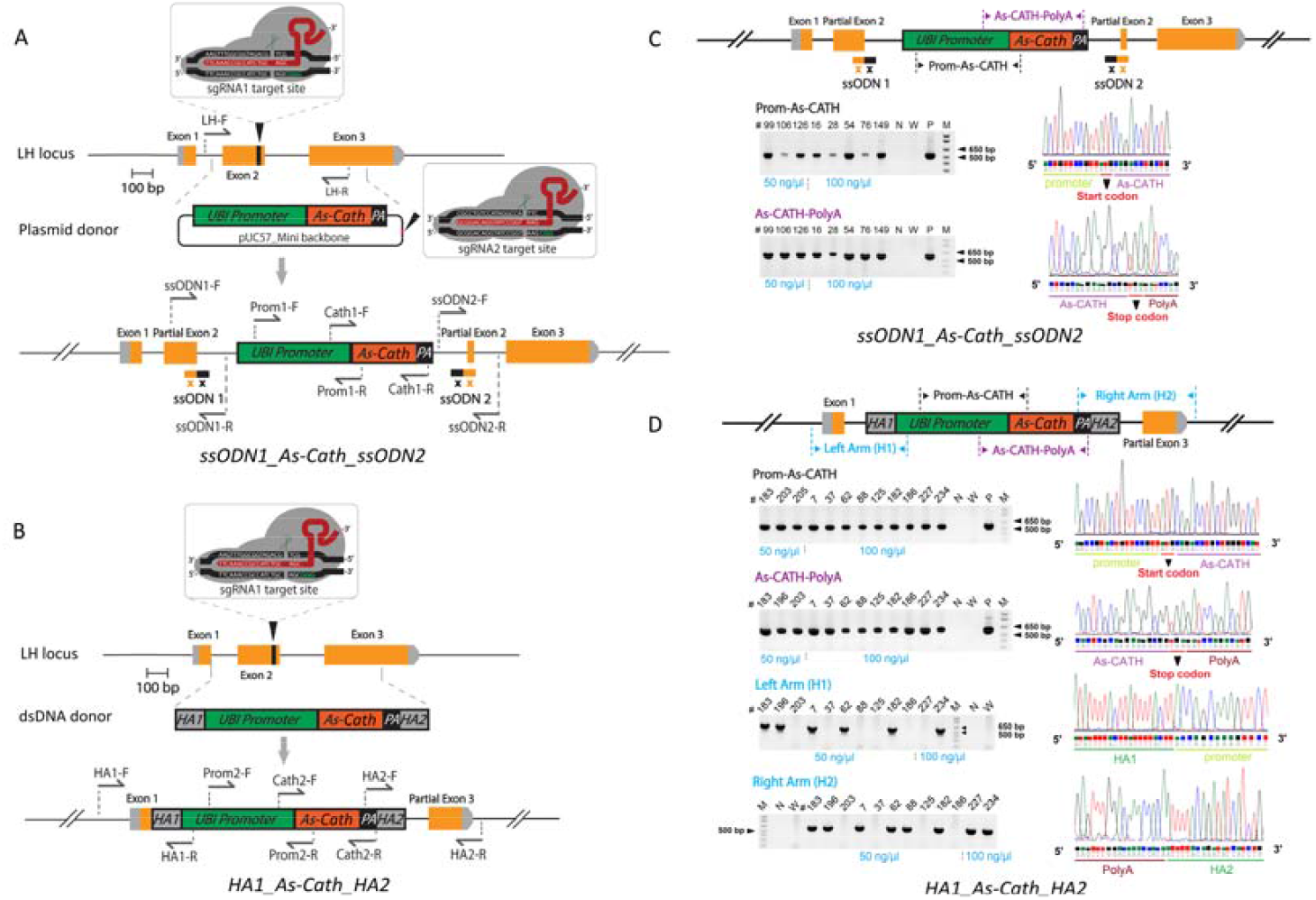
Single-stranded oligodeoxynucleotide (ssODN) and linear double-stranded DNA (dsDNA) with CRISPR/Cas9 mediating knock-in (KI) at the *luteinizing hormone* (*LH*) locus of channel catfish. **(A)** Schematic illustration of the insert-specific region for the cathelicidin gene from *Alligator sinensis* (*As-Cath*) KI via the two-hit two-oligo (2H2OP) system assisted by ssODNs at the *LH* locus, named as the ssODN1_As-Cath_ssODN2 construct. The structure of the *LH* gene’s exons is constructed by yellow bars, sgRNAs-targeted sites are indicated by black triangles, and the target sequences are detailed in rectangular boxes. The protospacer-adjacent motif (PAM) is highlighted in green. Primer sets are illustrated, showing the strategy to test *LH* mutation, ssODN1/ssODN2 junctions, the UBI promoter region and the insert-specific region of the *As-Cath* gene using PCR amplifications. **(B)** Schematic diagram of the *As-Cath* KI via the dsDNA system, named HA1_As-Cath_HA2 donor. Primers show the strategy to test the HA junctions, UBI promoter region, and *As-Cath* gene region. **(C)** TAE agarose gel of PCR amplicons showing off-target positive detection of the ssODN1_As-Cath_ssODN2 construct using 2H2OP method. The promoter region (Prom-As-CATH, 519 bp) and *As-Cath* region (As-CATH-PolyA, 591 bp) were illustrated with sequencing results. **(D)** TAE agarose gel of PCR amplicons showing on-target positive detection of the HA1_As-Cath_HA2 construct using dsDNA method. The targeted gene regions (Prom-As-CATH, 542 bp and As-CATH-PolyA, 597 bp) and the junctional regions (HA1, 573 bp and HA2, 598 bp) were determined with sequencing results. The numbers on the top of the gel images indicate the sample IDs of the fish. Lane N, negative control using water as template; Lane W, wild-type control (nCT); Lane P, positive (plasmid or dsDNA donor) control; Lane M, DNA marker (1 kb), 500 and 650-bp bands are highlighted with black triangles; 50 and 100 ng/μL show the different doses of donors: plasmid or dsDNA.

### 2.3. Design of donor DNA, sgRNA and CRISPR/Cas9 system

Gene-targeted KI can be engineered via HDR using the dsDNAs or ssODNs as donor templates. In the current study, we employed two CRISPR/Cas9-mediated systems to conduct targeted KI of the As-Cath fragment at the LH locus. For the first system, the CDS of the *As-Cath* gene was cloned into the pUC57_mini vector at the EcoRV enzyme digestion site to create the ssODN1_As-Cath_ssODN2 construct as a plasmid donor. Two sgRNAs (sgRNA1 and sgRNA2) were co-injected to operate as “scissors”, cutting the *LH* gene and linearizing the plasmid donor, respectively, and provided two short ssODNs to ligate the ends of both cut sites, labeled as the 2H2OP system (Fig. 1(A)). ssODN1 consists of 80 bp, of which the upstream 40 bp are derived from partial exon 2 of *LH* gene and the remaining 40 bp are homologous to pUC57_mini backbone. For ssODN2, the upstream 40 bp are from the pUC57_mini backbone, while downstream 40 bps come from a portion of exon 2 of the *LH* gene. The dsDNA donor was created by constructing the As-Cath CDS sequence flanked with two homology arms (HAs) of 300 bp derived from the *LH* gene of channel catfish on either side of the insert DNA, and we tagged the second construct as HA1_As-Cath_HA2. More specifically, 163 bp of HA1 (the left homology arm) are derived from the upstream of exon 2; 136 bp are identical to intron 1, and 1 bp originated from exon 1. HA2 (the right homology arm) contains 21 bps from exon 2’s downstream; 85 bps from intron 2 and 194 bps from upstream of exon 3 (Appendix A). Here, we used one sgRNA (sgRNA1) to cut the LH site in the channel catfish genomic DNA and provided a linear dsDNA as the donor template, and this system was labeled as dsDNA (Fig. 1(B)). For both constructs, the expression of the *As-Cath* gene was driven by the zebrafish ubiquitin (UBI) promoter [33]. The linear dsDNA, circular plasmid and ssODNs were synthesized by Genewiz (South Plainfield, NJ).

The sgRNAs were selected via the CRISPR design online tool (CRISPR Guide RNA Design Tool, Benching, https://zlab.bio/guide-design-resources) that targeted the *LH* gene of channel catfish and the donor plasmid. Candidate sgRNA sequences were compared to the whole genome of channel catfish via the Basic Local Alignment Search Tool to avoid cleavage of off-target sites. In addition, putative off-target sites were excluded using the online tool Cas-OFFinder (http://www.rgenome.net/cas-offinder/) [34]. Eventually, sgRNA1 for LH locus and sgRNA2 for donor plasmid were obtained. The Maxiscript T7 kit (Thermo Fisher Scientific, Waltham, MA) was used to generate sgRNAs *in vitro*, according to the instructions. Then purified sgRNAs were prepared using the RNA Clean and Concentrator Kit (Zymo Research, Irvine, CA). The concentration and quality of sgRNAs were detected with Nanodrop 2000 spectrophotometer (Thermo Fisher Scientific, Waltham, MA) and 1% agarose gel with 1 × tris-borate-EDTA (TBE) buffer, respectively. The synthetic sgRNAs were diluted to a concentration of ∼ 300 ng/μL and then divided into PCR tubes (2 μL/tube), and stored at −80 □ until use. The Cas9 protein powder was purchased from PNA BIO Inc. (Newbury Park, CA), and was diluted with DNase/RNase-free water to 50 ng/μL, keeping at −20 □ until use. Single guide RNAs and universal primer used in this study are listed in Table 1. Two different dosages of the donor DNA template and two control groups were set up: 50 ng/μL, 100 ng/μL, sham-injected control (iCT, only the 10% phenol red solution was injected) and non-injected control (nCT, no injection) for each KI system.

**Table 1.**
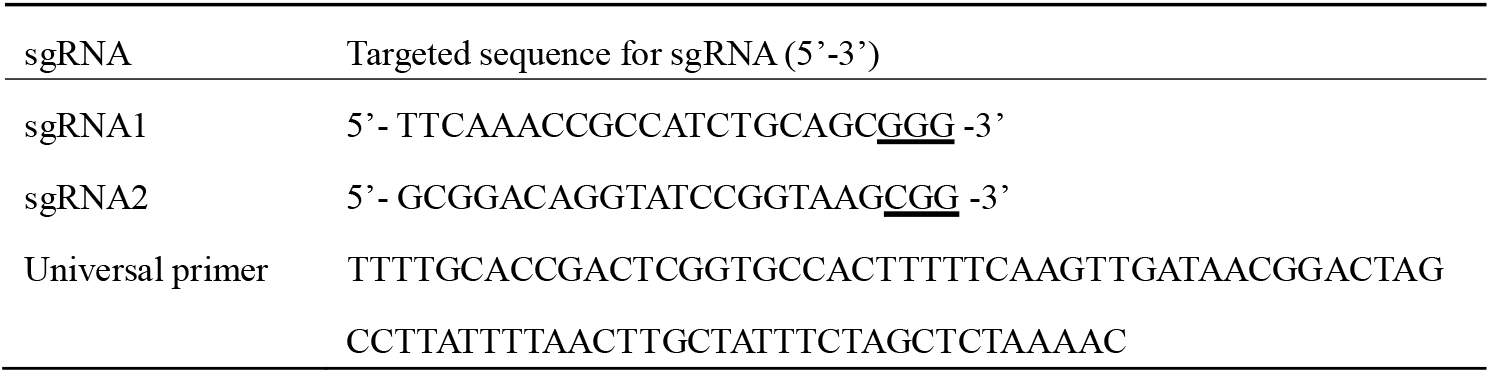
Target sequences of sgRNAs and the universal primer used in the present study. Underlined sequences represent the protospacer adjacent motif.

### 2.4. Transgenic fish production and rearing

Mature channel catfish females and males were paired for artificial spawning according to Elaswad et al. [35] with some modifications. Briefly, we selected individuals weighing more than 1.5 kilograms for spawning. Female channel catfish were implanted with 75 μg/kg of luteinizing hormone-releasing hormone analog (LHRHa) to induce ovulation, then eggs were gently stripped in a 20-cm greased spawning pan. Mature males were euthanized; testes were collected, rinsed, weighed and crushed; and sperm were prepared in 0.9 % saline solution (g:v = 1:10). Two milliliters of sperm solution were added to approximately 300 eggs and gently mixed. After a one-minute mixing, sufficient pond water was added to the eggs to activate the sperm, then the sperm/egg mixture was gently swirled for 30 s. More water was added and the embryos were kept in a single layer in the pan, and the embryos were allowed to harden for 15 min before microinjection.

The CRISPR/Cas9 system used for KI microinjections was combined with Cas9 protein, sgRNA and donor template in the ratio of 2:1:1, including one component of phenol red as an indicator. For the ssODN1_As-Cath_ssODN2 construct (2H2OP system), 8 μL of Cas9 protein (50 ng/μL), 2 μL of sgRNA1/sgRNA2 (300 ng/μL), 2 μL of donor plasmid (50 ng/μL, 100 ng/μL), 2 μL of ssODN1/ssODN2 (50 ng/μL, 100 ng/μL) and 2 μL of phenol red solution were mixed for microinjection (Total 8 + 2 + 2 + 2 + 2 + 2 + 2 = 20 μL). With respect to the HA1_As-Cath_HA2 construct (dsDNA system), 4 μL of Cas9 protein (50 ng/μL), 2 μL of sgRNA1 (300 ng/μL), 2 μL of donor dsDNA (50 ng/μL, 100 ng/μL), 2 μL of phenol red and 10 μL of DNase-free water were mixed to bring it up to 20 μL in total. For each mixture of the CRISPR/Cas9 system, we mixed Cas9 protein and sgRNA first and incubated them on ice for 10 min, then the donor templates were supplemented. For the iCT group, we only injected phenol red (diluted with 0.9 % saline). The mixed solution for each treatment was microinjected into one-cell stage embryos as previously described [36]. Every 6 μL of the mixture was loaded into a 1.0 mm OD borosilicate glass capillary that was pulled into a needle by a vertical needle puller (David Kopf Instruments, Tujunga, CA), and injected into 600 embryos. We injected 1,000 embryos dividing them into 5 random replicates for each treatment, and another 200 embryos with 3 replicates were prepared for each control group, respectively. All these embryos were from the same parents, and the microinjection was terminated after 90 min post-fertilization.

All injected and control embryos were transferred into 10-L tubs filled with 7-L Holtfreter’s solution (59 mmol NaCl, 2.4 mmol NaHCO_3_, 1.67 mmol MgSO_4_, 0.76 mmol CaCl_2_, 0.67 mmol KCl) [37] and 10−12 ppm doxycycline for hatching immediately after microinjection. All tubs were placed in the same flow-through hatching trough and a heater was put upstream of the trough to ensure that the water temperature was 26 −28 □ while dissolved oxygen levels were > 5 ppm via continuous aeration with airstones. Holtfreter’s solution was replaced twice per day and dead embryos/fry were collected and recorded daily during hatching to analyze hatchability, fry survival rate and genotype. The hatched fry were transferred to a Holtfreter’s solution without doxycycline and fed with live *Artemia* nauplii four times per day. After one week of culture in tubs, all fry from each treatment were stocked separately into 60 L aquaria (120 fish/tank) in a recirculating system for growth experiments. Feed pellet size was adjusted according to the size of the fish’s mouth as the fish grew. In detail, fry in tanks fed with Purina® AquaMax® powdered feed (50% crude protein, 17% crude fat, 3% crude fiber, and 12% ash) four times per day for two months. Then fingerlings were fed with Aquaxcel WW Fish Starter 4512 (45% crude protein, 12% crude fat, 3% crude fiber, and 1% phosphorus) twice a day for two months. Juvenile fish were fed with WW 4010 Transition feed (40% crude protein, 10% crude fat, 4% crude fiber, and 1% phosphorus) once a day [14]. All fish were fed to satiation.

### 2.5. Integration analysis and mutation detection

After a 4-month culture, all fingerlings (20−40 g) were pit-tagged (Biomark Inc., Boise, Idaho, USA) to distinguish each individual, the fish from different treatments were then mixed together and randomly dispersed into two circular tanks (1,200 L volume filled with ∼800 L of water) with the same density (120 fish/tank) for growth comparison monthly. Meanwhile, the pelvic fin clip and barbel were taken from anesthetized fish for DNA extraction and genotypic identification. During this phase, all fish received WW 4010 Transition feed once a day to satiation. Different genotyping strategies were involved for these two constructs: ssODN1_As-Cath_ssODN2, the CDS region of As-Cath was amplified to confirm gene insertion using primers Cath1-F/R (forward and reverse), and the promoter region was amplified via primers Prom1-F/R. As for the junctions, ssODN1 and ssODN2 regions were amplified using primers ssODN1-F/R and ssODN2-F/R to determine whether it was a target-site insertion. With respect to the HA1_As-Cath_HA2 construct, the As-Cath and promoter regions were detected using primers Cath2-F/R and Prom2-F/R, respectively. Then the left HA and right HA junctions were amplified via primers HA1-F/R and HA2-F/R. Primers were designed using the online software Primer3Plus (http://www.bioinformatics.nl/cgi-bin/primer3plus/primer3plus.cgi) and listed in Table S1 (Appendix B). PCR was performed in a 10-μL system and PCR products were resolved and visualized by running 1.0% agarose gel with 1 × tris-acetate-EDTA (TAE) buffer, and a bright band of each region with the corresponding length indicated an on-target positive (LH^−^_As-Cath^+^). Here, if we can determine that some individuals have been inserted with the As-Cath fragment, but we can not detect the junctional regions (HA-or ssODN-region), we then conclude them as potential off-target positives (LH^+^_As-Cath^+^).

With respect to the LH^+^_As-Cath^+^ fish, we selected 60 individuals to be tested for *LH* mutations. In this case, PCR was performed in a 20 μL-volume system using Expand High Fidelity^PLUS^ PCR System (Roche Diagnostics, Indianapolis, IN, USA) according to Elaswad et al. [35], and LH-F/R primers were used in both constructs. Then, the surveyor mutation detection assay was performed via Surveyor Mutation Detection Kit (Integrated DNA Technologies, IDT, Coralville, Iowa, USA) according to the detailed instructions [38]. A negative control reaction was included in the assay by using genomic DNA from the nCT group. Surveyor-digested DNA samples were electrophoresed for 1 hour in a 2% agarose gel using 1 × TBE buffer and compared to wild-type samples.

### 2.6. DNA sequencing

For the integrated As-Cath, promoter and junction sequences, PCR of positive samples was performed in a 50 μL-volume of system. Then the PCR products were purified using the QIAquick^R^ PCR Product Purification Kit (QIAGEN, Hilden, Germany) according to the manufacturer’s instructions. Before sequencing, all purified DNA samples were quantitated and identified using Nanodrop and by running 1.0% agarose gel. Primers Cath1-F/Cath2-F and Prom1-F/Prom-2F were used for sequencing of As-Cath and promoter regions for HA1_As-Cath_HA2 and ssODN1_As-Cath_ssODN2 constructs, respectively; primers HA1-F/HA2-F and ssODN1-F/ssODN2-F were used for sequencing of junctional regions for these two constructs, respectively.

Regarding LH mutations, we cloned the PCR products of putative mutant individuals using TOPO TA Cloning Kit (Invitrogen, Carlsbad, CA) before sequencing following the instructions with some modifications. Briefly, PCR was performed on each mutant individual that was previously identified with Surveyor assay using the primers LH-F/R for the next cloning steps. In addition, the DNA of three wild-type individuals from the nCT group was prepared using the same primers and procedures, then combined into one reaction and cloned as a wild-type control for sequencing. After cloning, we transformed the pCR™4-TOPO vector containing the PCR products into One Shot TOP10 Electrocomp™ *E. coli* (Invitrogen, Carlsbad, CA) as previously described [35]. Then 15 single colonies were randomly picked up to perform Colony PCR, and LH-F primer was used for the sequencing of LH mutant samples.

### 2.7. Determination of mosaicism and transgene expression

Five 12-month-old on-target positive fish and five sham-injected control fish were randomly chosen and sacrificed. Fourteen tissues, including skin, liver, kidney, spleen, blood, intestine, gill, stomach, fin, barbel, muscle, eye, brain and gonad of each individual were collected in 1.5 mL tubes and immediately transferred into liquid nitrogen for DNA and RNA isolation. PCR and quantitative real-time PCR (qRT-PCR) were conducted to detect the *As-Cath* gene’s potential mosaicism and mRNA level. Total RNAs were isolated from various tissues using TRIzol reagent (Thermo Fisher Scientific) and were reverse transcribed to cDNA using iScript™ Synthesis Kit (Bio-Rad, Hercules, CA) following the manufacture protocols.

qRT-PCR was performed on a C1000 Thermal Cycler using SsoFast™ EvaGreen Supermix kit (Bio-Rad, Hercules, CA) according to the instructions. Concentrations of the cDNA products were diluted to 250 ng/μL, and 1 μL template was used in a 10 μL PCR reaction volume. The mRNA level of 18S rRNA was used as an internal control, and the detailed qRT-PCR procedure was set up according to Coogan et al. [39]. The primers (Cath_RT-F and Cath_RT-R) used for qRT-PCR are listed in Table S1 (Appendix B). The CFX Manager Software (version 1.6, Bio-Rad) was used to collect the raw crossing-point (C_t_) values. The expression level of a target gene to the 18S rRNA gene from transgenic fish against non-transgenic sibling fish was converted to fold differences. Each sample was analyzed in triplicate using the formula 2^(−ΔΔCT)^, which sets the zero expression of the non-transgenic full-siblings to 1× for comparison.

### 2.8. Reproductive evaluation and restoration of parental KI fish

All P_0_ fish were stocked into a 0.04-ha earthen pond at Fish Genetics at Auburn University for growth and maturation. At the age of two years, some P_0_ individuals are expected to reach sexual maturity [40]. To evaluate the reproduction of two-year-old KI founders, on-target positive (LH^−^_As-Cath^+^), off-target positive (LH^+^_As-Cath^+^), and wild-type (WT) fish were selected to conduct a three-round mating experiment. Firstly, 3 pairs of WT, 6 pairs of LH^−^_As-Cath^+^, and 4 pairs of LH^+^_As-Cath^+^ mature parents were randomly placed into 13 tanks (60 × 45 × 30 cm^3^) for a two-week natural spawning to evaluate the spawnability of each genotype, and egg masses were collected from the spawnable parents. Then we primed the males with a 50 μg/kg LHRHa implant and 1600 IU/kg human chorionic gonadotropin (HCG) in the unspawned groups with a one-week observation to determine if LH^−^_As-Cath^+^ females were fertile. After this period, we recruited 6 more pairs of LH^−^_As-Cath^+^ fish to perform a 3 × 4 factorial design with 3 dosages of a combination of HCG and LHRHa implant (1200 IU/kg HCG + 50 μg/kg LHRHa, 1600 IU/kg HCG + 50 μg/kg LHRHa, 2000 IU/kg HCG + 50 μg/kg LHRHa) and 0.85% NaCl injected control group to assess the effects of hormone therapy. A 30-g egg mass for each genotype with 3 replicates was collected to calculate the fecundity (eggs/kg body weight [BW]). The masses were then transferred into tubs for hatchability and fry survival determination. Fish were fed ad libitum throughout the experiment.

### 2.9. Generation and genotype analysis for F_1_ fish

All the fry were separated into 60 L tanks by different genotypes. After 4 months of culture, fin clips and barbels were collected for DNA extraction from 60 F_1_ individuals of each genotype except the control groups. The same culture and genotyping procedures as described above were applied to the F_1_ generation.

### 2.10. Experimental challenge with Flavobacterium covae and Edwardsiella ictaluri

Gene-edited channel catfish were cultured in 60 L aquariums in the greenhouse of the Fish Genetics Laboratory at Auburn University (approved by AU-IACUC). To determine the resistance against pathogens, both P_0_ and F_1_ fish were challenged by *F. covae* and *E. ictaluri*.

#### *F. covae* challenge

Healthy P_0_ fingerlings with body weight 150.62 ± 4.24 g (mean ± SEM), including four genotypes (15 fish/genotype): LH^−^_As-Cath^+^, LH^+^_As-Cath^+^, negative LH^+^_As-Cath^−^ (negative fish without As-Cath insertion or LH mutation) and WT were mixed and acclimated in one hatching trough for five days and then transferred to a 1,800-L tank in the challenge room for acclimation for another 24 to 48 h prior to bacterial infections. All fish were randomly/equally separated into two 60-L buckets (30 L water). Briefly, a revived *F. covae* isolate (strain ALG-00-530) on modified Shieh agar (MSA) was inoculated into multiple cultures of 12 mL of modified Shieh broth (MSB) in 50-mL sterile flasks and grown in a shaker incubator at 150 rpm for 12 hours at 28°C. These cultures were then expanded into 200 mL cultures (5 mL additions) in 500 mL flasks and grown for another 12 h. The optical density was adjusted to OD_540_ = 0.731 and then spread plate dilutions were performed to determine the final inoculum concentration. One hundred microliters of each inoculum were serially diluted and spread onto MSA agar plates in duplicate and incubated at 28 °C for 48 h to quantify the concentration of the inoculum. Two flasks containing 325 mL of inocula (4.55 × 10^8^ CFU/mL) were immediately added to two 60 L buckets with fish following preparation, respectively. Then the fish were immersed statically in buckets for 1.5 hours at ∼28 °C (immersion dose: 2.46 × 10^6^ CFU/mL); afterward, all fish were gently moved back into the 1,800-L tank containing 1,000-L water and water flow was resumed. Meanwhile, a mock-challenged tank was used as the control but incorporated another 40 fish in 30 L of rearing water for 1.5 hours with sterile modified Shieh broth (325 mL) instead of the bacterial culture. With respect to the challenge of F_1_ fry (3.15 ± 0.24 g), four families of F_1_ fry (45 fish/family): LH^−^_As-Cath^+^, LH^+^_As-Cath^+^, LH^+^_As-Cath^−^ and WT were selected, and each family was randomly divided into three replicates with 15 fish per basket. The same challenge procedure and strain of *F. covae* with a dose of 4.75 × 10^8^ CFU/mL (immersion dose: 2.57 × 10^6^ CFU/mL) were implanted for the F_1_ generation.

#### *E. ictaluri* challenge

Sixty P_0_ fish (142.62 ± 3.72 g) including the above four genotypes, were prepared for the *E. ictaluri* challenge. *E. ictaluri* (S97-773) was provided by the USDA-ARS, Aquatic Animal Health Research Unit, Auburn, AL. The detailed procedures of the *E. ictaluri* challenge were performed according to Simora et al. [30] with some modifications. Briefly, 1 mL of frozen glycerol stock of *E. ictaluri* was inoculated into 20-mL brain–heart infusion broth (BHIB; Hardy Diagnostics) at 26°C in a shaker incubator at 180 rpm for 24 hours. And then bacteria were subcultured into 1-L BHIB for another 24 hours at the same condition until the cell density reached ∼1×10^8^ CFU/mL based on the OD_600_ value. All 60 P_0_ individuals were transferred into one 1,800-L tank for the challenge. Before starting *E. ictaluri* infection, water was lowered to a total of 100 L, then one liter of *E. ictaluri* suspension containing 3.2×10^8^ CFU/mL cells was added to the tank resulting in a final immersion dose of 3.2×10^6^ CFU/mL. Fish were immersed statically for 2 hours with aeration > 5 ppm, then water was restored. In addition to infected groups, one control tank containing 30 fish received only BHIB as a mock-challenged group. With respect to the challenge of F_1_ fingerlings (54.27 ± 1.49 g), a total of four genotypes containing 60 fish were selected, and the same challenge procedure and strain of *E. ictaluri* with a dose of 2.8 × 10^8^ CFU/mL (immersion dose: 2.8 × 10^6^ CFU/mL) were implanted for the F_1_ generation.

During the first 72 h of the experiment, we checked for mortality every four hours and then three times daily. Challenged fish were continuously monitored for 10 days for external clinical signs of *F. covae*/*E. ictaluri* and confirmation of bacteria colony growth by isolating bacteria from the kidney and liver to determine the cause of death, and dead individuals were recorded over time.

### 2.11. Statistical analysis

Spawnability, hatchability, fecundity, fry survival rate, and growth data were analyzed using one-way ANOVA/Tukey’s multiple comparisons test to determine the mean differences among treatments. To compare the KI efficiency of different groups, one-way ANOVA/Tukey’s multiple comparisons and odds ratio (OR) (Table S3 in Appendix B) were adopted. The survival curves of challenge experiments from different genotypes were compared by the Kaplan-Meier plots followed by Log-rank (Mantel-Cox) test. All statistical analysis was achieved via GraphPad Prism 9.4.1 (GraphPad Software, LLC). Gene expression between transgenic and non-transgenic fish was analyzed with an unpaired Student’s two-sample *t*-test. Statistical significance was set at *P* < 0.05, and all data were presented as the mean ± standard error (SEM).

## 3. Results

### 3.1. Targeted KI of As-Cath gene into the LH locus

Both the 2H2OP and dsDNA systems can induce As-Cath-integrated catfish lines with high integrated ratios, but the 2H2OP system had significant off-target effects (Fig. 1(CD), Fig. S1-S4 in Appendix B). More specifically, the 2H2OP system containing 50 ng/μL of donors (2H2OP50) showed the highest KI efficiency at 27.61% (37/134), followed by the groups 2H2OP100 (17.76%, 27/152), dsDNA50 (12.21%, 26/213) and dsDNA100 (10.25%, 25/244) (Table S2 in Appendix B). Although the 2H2OP50 group can introduce the highest KI efficiency (*P* < 0.01) (Fig. 2(A)), and 2H2OP system or 50 ng/μL of donors bring a significantly higher KI efficiency than the dsDNA method (*P* = 0.0001) or 100 ng/μL of donors (*P* = 0.00469) (Fig. 2(BC)). However, the dsDNA with 50 ng/μL donors demonstrated the highest on-target KI efficiency (10.80%, 23/213) compared to other treatments (*P* < 0.01) (Fig. 2(D)). In contrast, only one on-target KI case was observed in the 2H2OP system, which was significantly lower than that in the dsDNA (*P* < 0.0001) (Fig. 2(E)). Although different dosages of donors exhibited a significant effect on the total KI efficiency, our results indicated that this difference was not significant in the on-target KI (*P* = 0.3577) (Fig. 2(F)).

**Fig. 2.**
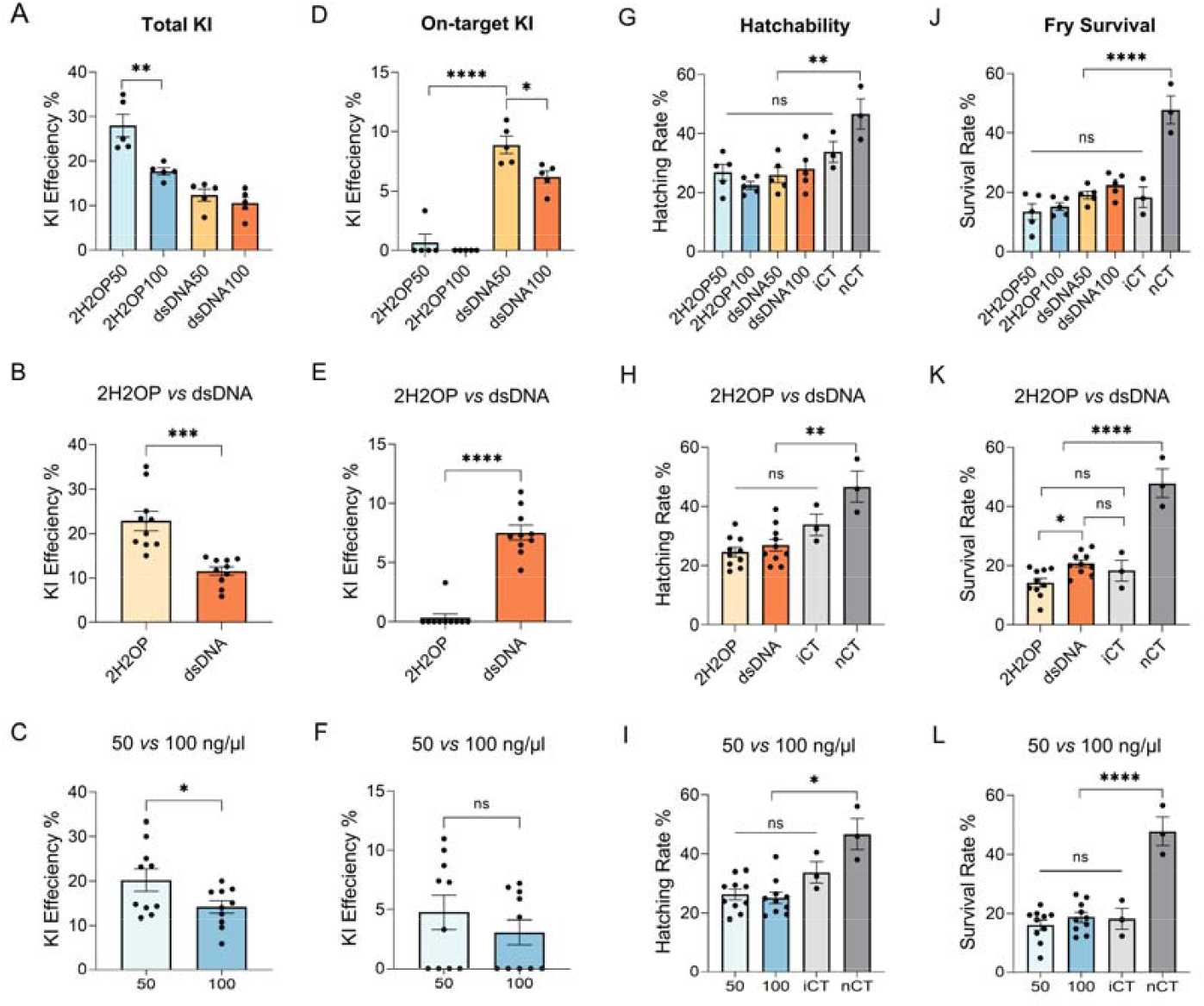
Effects of different CRISPR/Cas9-mediated systems (2H2OP *vs* dsDNA) with various dosages of donors (50 *vs* 100 ng/μL) on the knock-in (KI) efficiency, hatchability and fry survival rate. **(A)** Total KI efficiency of different CRISPR/Cas9-mediated systems and dosage combinations. **(B, C)** Comparison of total KI efficiency for different systems or dosages of donors. **(D)** On-target KI efficiency of different CRISPR/Cas9-mediated systems and dosage combinations. **(E, F)** Comparison of on-target KI efficiency of different systems or dosages. **(G)** Effect of different CRISPR/Cas9-mediated systems and dosage combinations on hatchability. **(H, I)** Comparison of the hatchability for different systems or dosages. **(J)** Effect of different CRISPR/Cas9-mediated systems and dosage combinations on fry survival. **(K, L)** Comparison of the fry survival rate for different systems or dosages. iCT, sham-injected control; nCT, non-injected control; 2H2OP(50/100), the CRISPR/Ca9-medicated system with ssODN1_As-Cath_ssODN2 construct (with a pUC57_mini plasmid and ssODN donor as 50/100 ng/μL); dsDNA(50/100), the CRISPR/Ca9-medicated system with HA1_As-Cath_HA2 donor DNA (with a dsDNA donor as 50/100 ng/μL); * = *P* < 0.05; ** = *P* < 0.01; *** = *P* < 0.001; **** = *P* < 0.0001; ns = not significant, by unpaired student’s *t*-test or one-way ANOVA.

According to the odds ratio, the 2H2OP system and low dosage tended to bear a higher total integrated rate which was 2.30 and 1.47 times than that of the dsDNA (OR = 2.30 for 2H2OP *vs* dsDNA) and high dosage (OR = 1.47 for 50 *vs* 100 ng/μL), respectively. Nonetheless, dsDNA had an overwhelming surpriority in on-target integration, which was more than 20 times greater than that in the 2H2OP system (OR = 26.70) (Table S3 in Appendix B). Taken together, the dsDNA system accompanied by a dosage of 50 ng/μL of donors tends to yield the highest on-target KI efficiency in our current study.

Given the non-As-Cath-integrated fish, we did detect individuals with only the LH mutation. Specifically, 5.56% (3/54), 6.67% (4/60), 3.33% (2/60), and 3.33% (2/60) of fish with *LH* deficiency in the 2H2OP50, 2H2OP100, dsDNA50 and dsDNA100 groups, respectively, were detected by the Surveyor mutation test (Table S2 in Appendix B). The sequencing results revealed that 2, 2, 1 and 3 types of mutations in 4 *LH*-mutant individuals from the 2H2OP100 group (Fig. S5 in Appendix B).

### 3.2. Effects of the dosage and CRISPR/Cas9 system

Different donor dosages and CRISPR/Cas9-mediated systems exhibited toxicity to fish embryos by decreasing the hatchability and fry survival rate. Although there were no significant differences in hatching rates among these four CRISPR/Cas9-mediated injected groups compared to the iCT group (*P* = 0.1630), the hatching rate was lower than the nCT group (*P* < 0.01) (Fig. 2(G)). Moreover, the lethality of embryos was consistent across different donor dosages (50 *vs* 100 ng/μL) (*P* = 0.1080) or CRISPR/Cas9-mediated systems (2H2OP *vs* dsDNA) (*P* = 0.0796), which was significantly higher than that in the nCT group (Fig. 2(HI)). For the fry survival, the survival rate of the microinjection group was significantly lower compared with the nCT group (*P* < 0.0001) (Fig. 2(J)). In addition, the dsDNA system induced a higher survival rate of fry (*P* = 0.0031) (Fig. 2(K)) than the 2H2OP system. Still, donor dosages showed no significant differences in fry survival after hatching (*P* = 0.2923) (Fig. 2(L)).

### 3.3. Mosaicism and As-Cath expression

PCR and RT-PCR were used to detect the *As-Cath* transgene and its expression of different tissues in on-target positive fish. The results revealed that three of the five LH^−^_As-Cath^+^ fish showed the expression of the *As-Cath* in all 14 sampled tissues (skin, liver, kidney, spleen, blood, intestine, gill, stomach, fin, barbel, muscle, eye, brain and gonad) (Fig. 3(AB)), but one of them had expression observed in 11 tissues (except barbel, muscle and gill) and another one in 8 tissues (skin, liver, blood, intestine, gill, barbel, muscle and gonad) (Fig. S6 in Appendix B), suggesting mosaicism in the on-target positive individuals. We found that the expression of *As-Cath* was detected even without pathogenic infections for the three on-target positive individuals. The three highest mRNA levels were determined in the kidney (28.91 fold changed), skin (24.30 fold), and gill (8.445 fold), followed by the muscle (7.430 fold), spleen (6.047 fold) and barbel (4.808 fold). However, the eye (1.327 fold), intestine (1.589 fold), and fin (1.608 fold) had the lowest expression compared to other tissues (Fig. 3(C)).

**Fig. 3.**
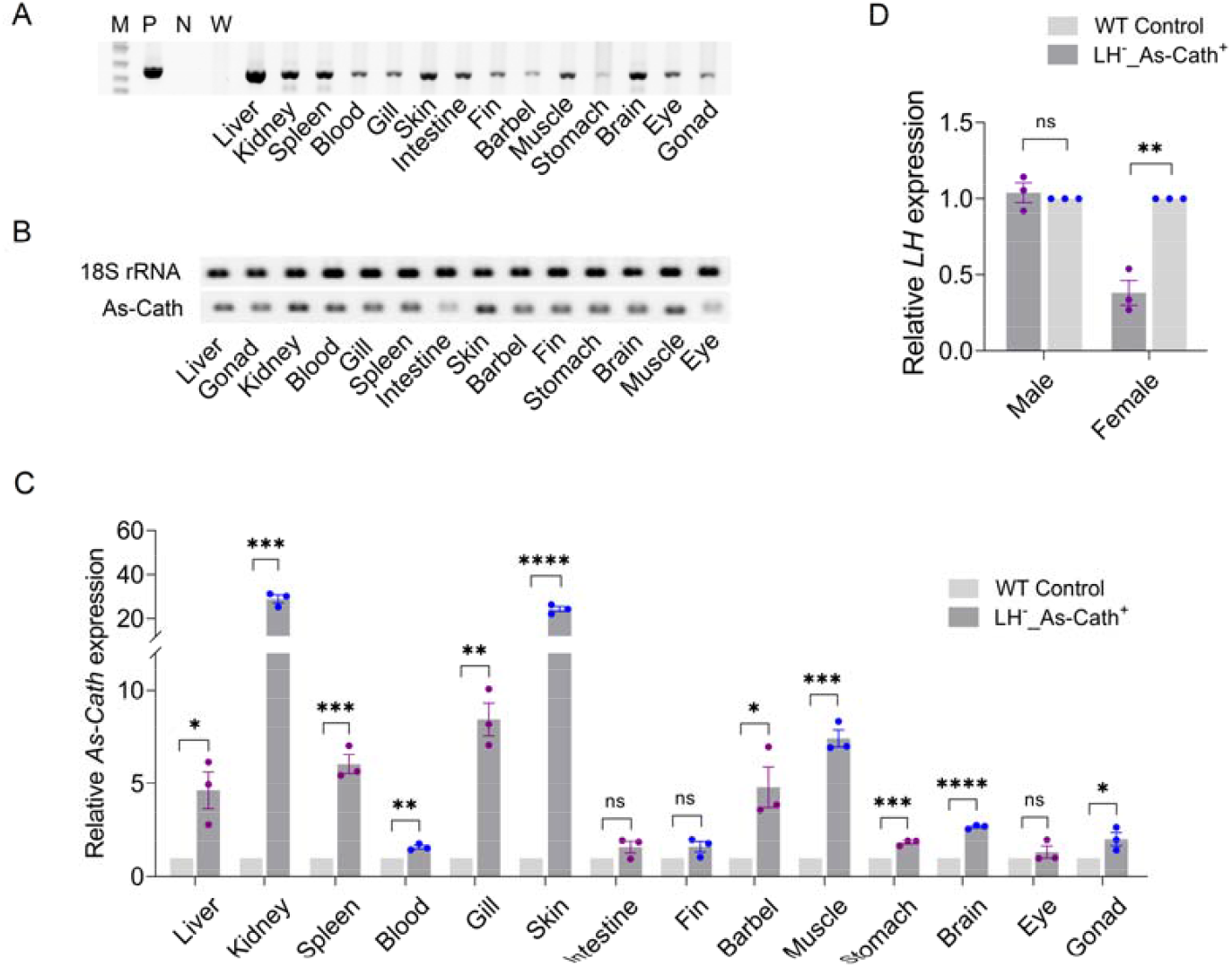
Mosaicism detection and the expression of the cathelicidin gene from *Alligator sinensis* (*As-Cath*) in the LH^−^_As-Cath^+^ fish line. **(A)** PCR amplicons show the *As-Cath* region in 14 tissues from one representative LH^−^_As-Cath^+^ fish. **(B)** The agarose gel electrophoresis showed the *As-Cath* gene expression in various tissues of P_0_ transgenic channel catfish, *Ictalurus punctatus*. **(C)** Relative *As-Cath* gene expression of different tissues from RT-PCR analyses. **(D)** Relative *LH* gene expression of gonads from LH^−^_As-Cath^+^ males and females. Expression levels were calibrated against corresponding tissues from sibling wild-type fish, and three individuals were employed for each genotype. Lane M, DNA marker (1 kb); Lane P, positive (plasmid or dsDNA donor) control; Lane N, water negative control; Lane W, wild-type control (nCT); * = *P* < 0.05; ** = *P* < 0.01; *** = *P* < 0.001; **** = *P* < 0.0001; ns = not significant, by unpaired student’s *t*-test or one-way ANOVA.

In addition, compared to the WT individuals, the mRNA level of *LH* in gonads was down-regulated in LH^−^_As-Cath^+^ females at the age of one year (*P* = 0.0016), but there was no significant difference in that of males (*P* = 0.5817) (Fig. 3(D)).

### 3.4. Reproductive sterility and restoration of reproduction

A three-round mating experiment determined the promise for complete control of channel catfish reproduction (Fig. 4(A)). Our outcomes revealed that three pairs of WT (100%, 7927 eggs/BW) and two pairs of LH^+^_As-Cath^+^ fish (50%, 8952 eggs/BW) were spawned respectively during the first two-week natural mating, but no spawn was observed in the LH^−^_As-Cath^+^ pairs (0%). Compared to the LH^−^_As-Cath^+^ pairs, WT and LH^+^_As-Cath^+^ fish had higher spawnability under natural pairing conditions (*P* = 0.0148 and *P* = 0.1743). In addition, the LH^+^_As-Cath^+^ pairs did not show a significant difference in spawnability compared to the WT pairs (*P* = 0.2143) (Fig. 4(B)).

**Fig. 4.**
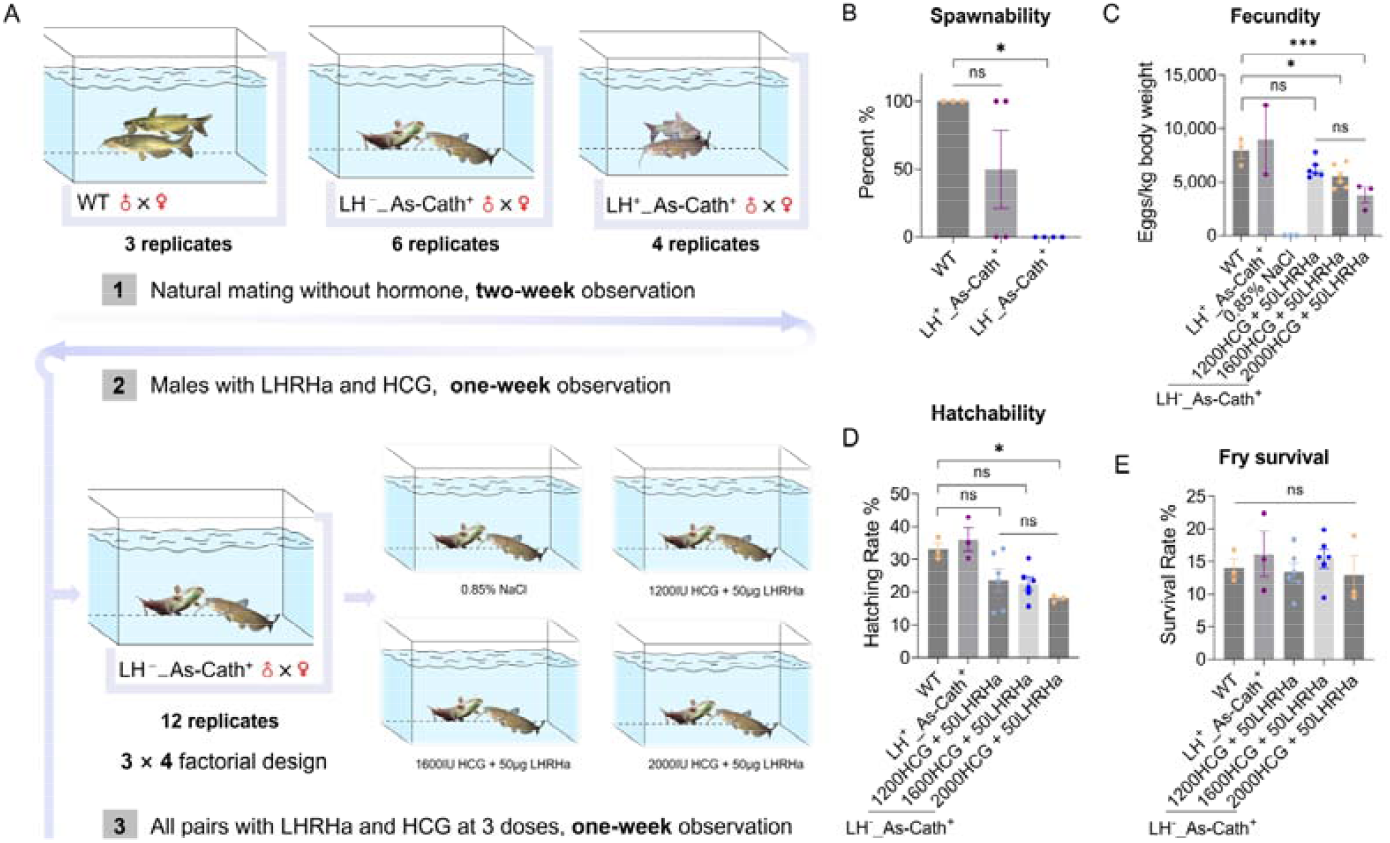
Reproductive determination and restoration of the As-Cath-integrated fish lines. **(A)** A three-round design of the reproduction experiment. Three genotypes of P_0_ founders: WT, LH^−^_As-Cath^+^, and LH^+^_As-Cath^+^ fish were involved. First round, 3, 6 and 4 pairs as replicates for each genotype were set up randomly in 13 tanks for mating without hormone treatments, and a two-week observation was adopted. Second round, moved out spawned pairs and primed un-mated males with a 50 μg/kg LHRHa implant and 1600 IU/kg HCG to determine the reproduction of LH^−^_As-Cath^+^ females, observing for one week. Third round, 12 pairs of LH^−^_As-Cath^+^ fish were complemented and re-paired and treated with three doses of LHRHa and HCG in a 3 × 4 factorial design for one week. **(B)** Detection of spawnability for LH^−^_As-Cath^+^ fish during natural mating. **(C, D, E)** Potential effects of different hormone treatments on the fecundity and hatchability of P_0_ generation, and fry survival of F_1_ generation. LH, luteinizing hormone; LHRHa, luteinizing hormone-releasing hormone analogue; HCG, human chorionic gonadotropin; * = *P* < 0.05; ** = *P* < 0.01; ns = not significant, by unpaired student’s *t*-test or one-way ANOVA.

Furthermore, a one-week hormone priming (50 μg/kg LHRHa + 1600 IU/kg HCG) of the males did not stimulate LH^−^_As-Cath^+^ females to give eggs, indicating LH-deficient females blocked oocyte maturation and ovulation. However, our results discovered that a combination of LHRHa and HCG can effectively induce spawning for the LH^−^_As-Cath^+^ females when both males and females were primed. Specifically, two, two and one female gave eggs after 24 to 48 hours post-hormone injection from the 1200 IU (6213 eggs/BW), 1600 IU (5514 eggs/BW) and 2000 IU/kg (3778 eggs/BW) HCG group combined with 50 μg/kg LHRHa, respectively. These three treatments significantly improved the fecundity compared to 0.85 % NaCl injection (*P* < 0.0001). Additionally, the fecundity decreased with increasing hormone dosage, but the difference among these three hormone dosages was not significant (*P* = 0.0731). Nevertheless, the fecundity can be restored to a normal level when 1200 (*P* = 0.2627) or 1600 (*P* = 0.1983) IU/kg HCG combined with 50 μg/kg LHRHa was adopted (Fig. 4(C)). Compared with the WT and the other hormonal-therapy groups, the 2000 IU/kg HCG group significantly reduced the fecundity (3778 eggs/BW, *P* = 0.0494) and hatchability (18.01%, *P* = 0.0476) (Fig. 4(D)). Although different hormonal treatments had varying effects on fecundity and hatchability, they had no effects on fry survival at the early stage (*P* = 0.1018) (Fig. 4(E)).

### 3.5. F_1_ genotyping, growth comparison in P_0_ and F_1_

As mentioned above, three WT, two LH^+^_As-Cath^+^, and five LH^−^_As-Cath^+^ families were generated from our three-round mating experiment. However, genotype analysis determined that only one family in the LH^+^_As-Cath^+^ line (33.33%[10/30] integrated rate in the F_1_ offspring) and two families in the LH^−^_As-Cath^+^ line (40%[12/30] integrated rate in the F_1_ progeny of family 1 and 46.67%[14/30] integrated rate in the F_1_ offspring of family 2), respectively, had the *As-Cath* gene detectable in the F_1_ generation. These results further confirmed the existence of the mosaic phenomenon in the P_0_ founders.

To determine the effects of *LH* disruption and *As-Cath* integration on fish growth, we compared the BW over time of the P_0_ founders and the F_1_ progeny, respectively. The growth data suggested that the LH^−^_As-Cath^+^ individuals did not show superiority in terms of growth in the first nine months in the P_0_ generation. Nonetheless, P_0_ LH^−^_As-Cath^+^ fish exhibited the largest body gain (36.35 g) compared to other genotypes (25 g). Furthermore, significantly faster growth was demonstrated in the F_1_ generation of LH^−^_As-Cath^+^ after a three-month culture. Hence, our results indicated more immediate growth potential for the LH^−^_As-Cath^+^ fish than the WT fish (Table 2).

**Table 2.**
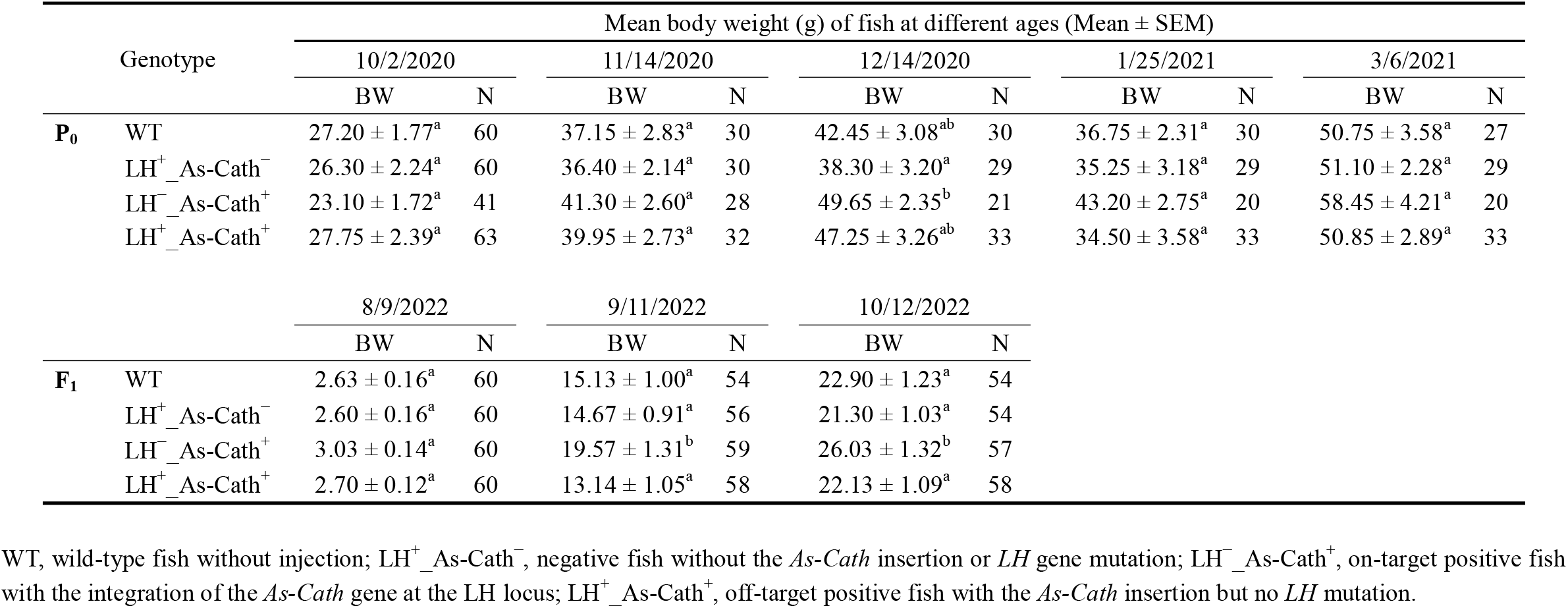
Mean monthly body weight (BW), sample size (N) over time of P_0_ and F_1_ As-Cath-integrated, negative and control channel catfish, *Ictalurus punctaus*. P_0_ founders were generated in June 2020, and F_1_ progeny were produced in June 2022. For both generations, four genotypes: WT, LH^+^_As-Cath^−^, LH^+^_As-Cath^+^, and LH^+^_As-Cath^+^ were included. Fish were kept separately in 60-L aquaria with the density of 2 fry/L until 4 months post hatch, then they were pit-tagged (10/2/2020) and transferred to a 1,200-L circular tank (∼800-L water) with a mix of these 4 genotypes (initial number of fish was 30, 30, 28 and 32) and fed daily to satiation. Differences in BW among these four genotypes were compared using one-way ANOVA followed by Tukey’s multiple comparisons test. Means with different letters as superscripts are significantly different (*P <* 0.05).

### 3.6. Enhanced resistance against fish pathogens

Enhanced resistance against *F. covae* and *E. ictaluri* of As-Cath-integrated fish was observed compared to WT/negative individuals from our challenge experiments in both P_0_ and F_1_ generations. According to *F. covae* challenge results, there was no significant difference in survival rate between the two types of controls (WT and LH^+^_As-Cath^−^) in both P_0_ (13.33% *vs* 20%, *P* = 0.8682) and F_1_ generation (26.67% *vs* 40%, *P* = 0.8955). However, LH^−^_As-Cath^+^ and LH^+^_As-Cath^+^ fish exhibited significantly improved survival post *F. covae* infection compared to the WT control group in both P_0_ founders (LH^−^_As-Cath^+^ *vs* WT: 73.33% *vs* 13.33%, *P* = 0.0016; LH^+^_As-Cath^+^ *vs* WT: 66.67% *vs* 13.33%, *P* = 0.0014) and F_1_ progeny (LH^−^_As-Cath^+^ *vs* WT: 86.67% *vs* 26.67%, *P* = 0.0010; LH^+^_As-Cath^+^ *vs* WT: 73.33% *vs* 26.67%, *P* = 0.0127). Additionally, on-target insertion of the *As-Cath* gene resulted in improved resistance against *F. covae* than in the off-target positives without statistically differing in both generations (73.33% *vs* 66.67%, *P* = 0.7726 for P_0_, and 86.67% *vs* 73.33%, *P* = 0.3613 for F_1_). Furthermore, our findings revealed that the F_1_ progeny was more resistant to *F. covae* than its P_0_ parents (Fig. 5(AB)).

**Fig. 5.**
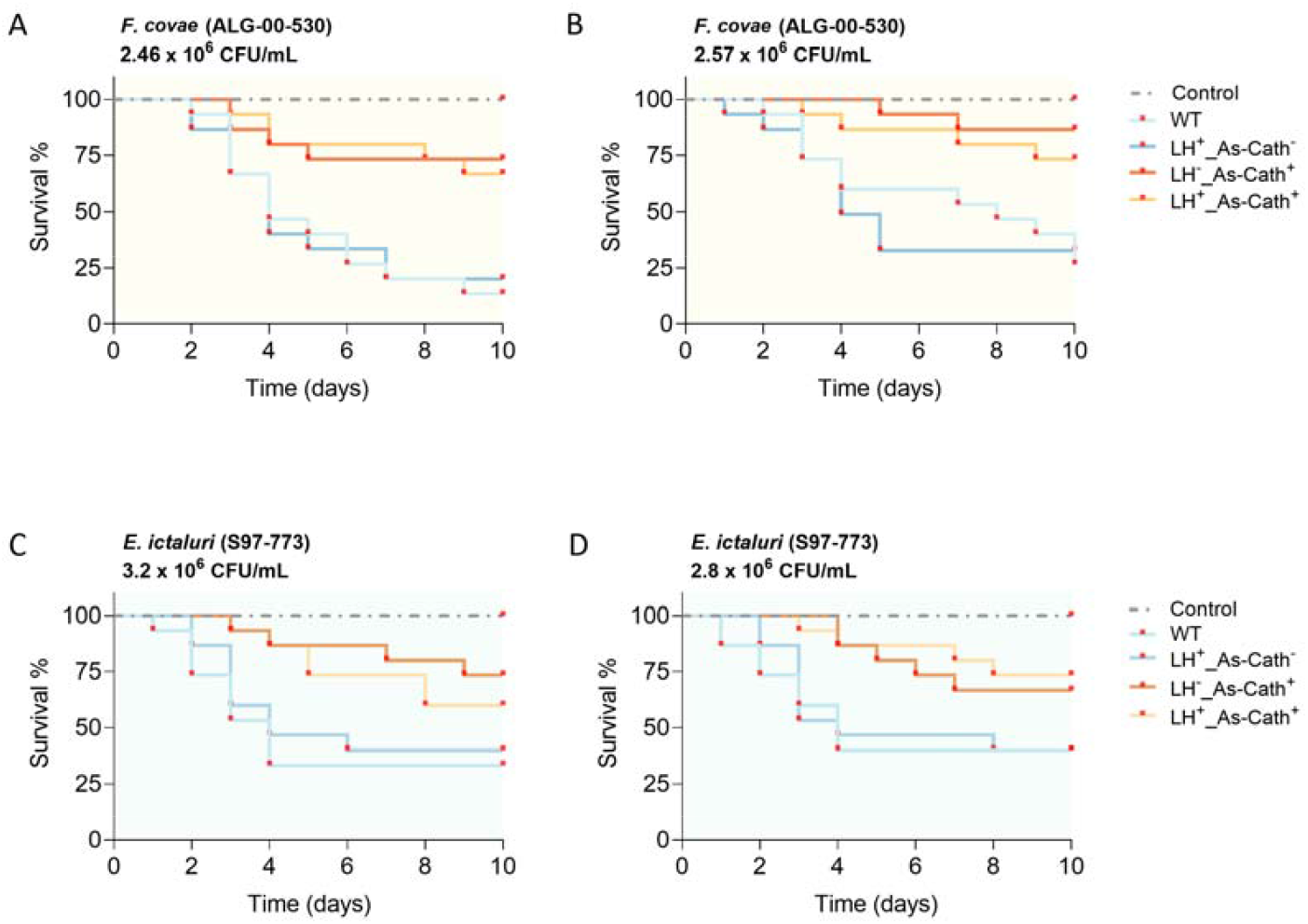
Kaplan-Meier plots of *As-Cath* integrated catfish against two fish bacterial pathogens. **(A, B)** Survival curves of P_0_ and F_1_ generations for a variety of genotypes infected by *Flavobacterium covae*, respectively. **(C, D)** Survival curves of P_0_ founders and F_1_ progeny for different genotypes infected by *Edwardsiella ictaluri*, respectively. In addition to these bacterial infection groups, one control group with medium immersion was implanted for each challenge experiment, and the immersion dose was presented in each figure. Comparison of different survival curves was determined by the Log-rank (Mantel-Cox) test. WT, wild-type, non-injected fish line; LH^+^_As-Cath^−^, negative fish line (micro-injected fish without *LH* mutation and *As-Cath* insertion); LH^−^_As-Cath^+^, on-target positive fish (*As-Cath* insertion was detected at *LH* locus); LH^+^_As-Cath^+^, off-target positive fish (*As-Cath* insertion was detected but not at *LH* locus).

Increased resistance to *E. ictaluri* was also observed in the P_0_ (LH^−^_As-Cath^+^ *vs* WT: 73.33% *vs* 33.33%, *P* = 0.0125; LH^+^_As-Cath^+^ *vs* WT: 60% *vs* 33.33%, *P* = 0.0427) and F_1_ generations (LH^−^_As-Cath^+^ *vs* WT: 66.67% *vs* 40%, *P* = 0.0558; LH^+^_As-Cath^+^ *vs* WT: 73.33% *vs* 40%, *P* = 0.0350), with results that were similar to those of the *F. covae* challenge. Overall, As-Cath-integrated individuals showed a significant improvement in the survival rate compared to the WT fish (66.67% *vs* 33.33%, *P* = 0.0381 for P_0_; 70% *vs* 40%, *P* = 0.0335 for F_1_). Nevertheless, there was no significant difference in LH^−^_As-Cath^+^ and LH^+^_As-Cath^+^ fish (73.33% *vs* 60%, *P* = 0.4566 for P_0_; 66.67% *vs* 73.33%, *P* = 0.6851 for F_1_) (Fig. 5(CD)).

## 4. Discussion

In contrast to the previous gene-editing oriented exclusively to the improvement of the desired traits, the present study took into account ways to lessen the potential impact of transgenic fish on the ecosystems and genetic biodiversity. Specifically, we successfully integrated an AMG into the reproduction-associated locus using different CRISPR/Cas9-mediated systems. We identified a suitable KI system for channel catfish to achieve boosted resistance against fish pathogens and reproductive control, reducing the reliance on antibiotics and anti-parasitics in aquaculture. The HA-mediated CRISPR/Cas9 system displayed a high integrated rate, low off-target events, and low toxicity. In addition, reproduction is entirely controllable and can only be restored to normal levels of fecundity with hormone therapy in the new fish line. In general, the insertion of the cathelicidin gene at the *LH* locus for enhanced resistance against infectious diseases and reproductive confinement to improve consumer-valued qualities and to promote the environmental friendliness of transgenic fish appears promising.

There have been several obstacles involved in the CRISPR/Cas9-mediated KI system when it is used in the embryos of non-model animals. In the history of genome editing, the initial CRISPR/Cas9 systems were proposed based on mammalian cells or embryos of the model animals. From model to non-model animals, there are several uncertainties, such as embryo size, developmental period, and the sensitivity to Cas9 protein that researchers have to optimize a fitted system when starting a new species’ genome editing. Yoshimi et al. [5] demonstrated that the ssODN-mediated end joining approach induced a high integrated rate of 17.6% (3/17) in rats when a short ssODN template was provided. Conversely, recent works indicated that ssODN-mediated KI could induce a high percentage (17.8%) of indel mutations in sheep [41]. In the current study, we used CRISPR/Cas9 systems mediated by ssODN and HA to create on-target KIs of the *As-Cath* gene at the *LH* locus. Although a high KI efficiency of 22.38% (64/286) was detected in the ssODN-mediated system, it caused a high off-target frequency (> 90%) in the channel catfish. Our results are in agreement with findings in zebrafish, which have illustrated that erroneous ssODN integration occurred when various template lengths were adopted [7]. These studies suggest that ssODN-mediated KI efficiency in fish models relies heavily on ssODN templates [42], and caution is warranted when employing ssODNs to create KI models.

Compared to the ssODN-mediated system, HA-assisted KI can achieve a 20–30% HDR-mediated knockin in human cells with various homogenous sequences [9,43]. In addition, Simora et al. [44] determined that HA-mediated CRISPR/Cas9 provided with a linear dsDNA donor displayed a total integrated rate of 29% at the non-coding region of channel catfish genome, which is drastically higher than that of this work (29% *vs* 11.16%[51/457]). We believe this difference in integration rate is due to the different sample sizes, unknown functions in the target regions (non-coding *vs LH* locus), efficiency of sgRNA and HA, and unpredictable genetic interaction; the larger sample size from our study could give more robust conclusions. These findings reveal that the HA-mediated system is more effective in the catfish species compared to the ssODN. The KI efficiency of HDR-induced CRISPR/Cas9 has been at a low level including in cell lines and model animals [5,7,9]. Fortunately, new CRISPR/Cas-mediated techniques are constantly being developed. For instance, the CRISPR/Cas12i-mediated system shows promise in multiplexed genome editing with high mutation rates in human T cells [45]. Additionally, Kelly et al. [46] established a CRISPR/Cas9 HITI system for the insertion of large DNA donors with high integrated efficiency of 36% in human 293T cells. Recently, a new approach named dCas9-SSAP demonstrated a high on-target KI efficiency (∼20%) knocking in long sequences in mammalian cells [47]. These new tools or systems are encouraging to be applied from model to non-model animals and could improve genome-editing efficiency.

Although we predicted and avoided possible off-target sites using the well-acknowledged software, the actual integration results showed the existence of off-target activities. This is mainly due to the failure of *in silico* prediction to predict *bona-fide* off-target sites *in vivo* [48,49]. Furthermore, the frequency of off-target events is higher *in vivo* of animal experiments than that of in cellular experiments *in vitro* [50]. The majority of published studies contend that the observed unintended mutations/insertions is one major concern in the application of the CRISPR/Cas9 system, which could confound the interpretation of findings [49,51,52]. However, although some reports claim that no detectable undesirable mutations/insertions from the genotypes or phenotypes have been revealed in mice and fish [44,53,54], the following underlying potentials could be noted: 1) Unaltered phenotypes may be observed since the off-target cleavage can occur in a non-coding region [55]. 2) The researchers tend to focus on the P_0_ founders with intended insertions rather than those harboring possible off-target mutations [56-57]. 3) Most published research using animal models does not use genome-wide methodologies for detecting off-target cases, which could conceal some infrequent off-target editing sites [50]. In the same case, with the exception of *LH* mutations, we did not conduct a thorough detection on all off-target individuals due to its being time-consuming and expensive. Nevertheless, this does not preclude us from keeping the non-analyzed off-target individuals as we will eventually genotype them in a genome-wide and unbiased way.

Genetic mosaicisms have been and will still be another obstacle to applying CRISPR/Cas9-mediated genome editing in practical applications. In this study, we failed to effectively obtain a 100% of individuals without mosaics. In essence, mosaicism from CRISPR/Cas9-genome-edited organisms is common in the case of fertilized egg-based editing, and mosaic animals have been observed in mice [58,59], rats [57] and zebrafish [60,61] with a variety of frequencies. CRISPR/Cas9 engineered mosaicisms bring undesired consequences, hindering the generation of homozygous positive offspring and prolonging the generation of homozygotes. We evaluated the *As-Cath* gene expression from five on-target positive P_0_ founders and found that one individual had no expression in the gonad. In our study, several mosaic events were determined in the germline, resulting in the inability to transfer the *As-Cath* gene to the offspring. Thus, we believe that mosaicism is also common and unavoidable in non-model fish. Although early sperm/testis or egg/ovary genotyping can be effective in avoiding the creation of undesirable offspring, it is challenging to access the germline DNA without sacrificing the parents. Of importance, we still maintain our mosaic populations for genotyping and phenotyping in the further F_2_ and F_3_ progeny until homozygous individuals are obtained. Future research could reduce mosaicisms by delivering CRISPR/Cas9 components to very early-stage zygotes [6]. Alternatively, the new strategies, i.e., *Easi*-CRISPR, C-CRISPR [6], CRISPR/Cas9 HITI [46] and dCas9-SSAP [47] could be used to prevent the induction of mosaic animals.

Regardless of the type of CRISPR/Cas9-mediated genome editing, microinjection always has irreversible effects on embryos, i.e., increased mortality and decreased hatchability from our current study. High embryonic deaths were observed from shame- and CRISPR/Cas9-mediated-microinjection in our study, revealing that major mortality occurs due to the injection of the yolk, while fewer impacts are from the DNA donors and reagents [44]. Although a high dosage resulted in a high embryonic mortality and lower hatching rate, it did not significantly reduce the fry survival rate compared to the injected-control group, which is in agreement with the findings from Elaswad et al. [35]. This may be because microinjection only has a detrimental effect on the yolk of the embryo. Still, this effect no longer affects the fry once the fertilized eggs have successfully hatched. Given the unavoidable physical lethality of embryos, off-target effects and mosaicisms, we recommend microinjection of ∼3000 fertilized eggs for non-model fish species in order to afford enough gene-edited fish for subsequent validation experiments.

To assess the pleiotropic effects, we compared the growth performance of the on-target/off-target As-Cath-integrated fish line with the WT population. Our findings demonstrated that off-target insertions did not exhibit growth depression or improvement in various families of P_0_ founders. Nonetheless, the preliminary data revealed the LH^−^_As-Cath^+^ fish had a greater gain in body weight compared to the WT individuals after a three-month culture in the tank, indicating that the growth differences are emerging in the F_1_ progeny. This variation may be due to heterozygous individuals lacking stable genetic traits, or off-target integrations in other regions concealing growth advantages in the P_0_ generation [50]. cfGnRH-deficient channel catfish did not show significant effects in growth and survival throughout a four-year culture compared to the WT fish [19]. However, potential pleiotropic effects could exist when the *LH* gene is replaced by the As-Cath in our cases. Therefore, P_0_ mosaic founders carrying the *As-Cath* gene should be used to produce F_1_, F_2_ and F_3_ homozygous families, and then the comparisons of the growth, survival rate, seinability and carcass traits could be performed to avail the enhanced performance of LH^−^_As-Cath^+^ fish line more transparent to farmers and the public in the future.

HDR-mediated KI is rarely applied in aquaculture due to the very low integration efficiency, but most of the traits were achieved by NHEJ-mediated KO [17,18]. In addition, few studies proved that gene-mutants can induce disease-resistant fish lines via KO to date [13]. By contrast, the integration of AMG is encouraging to improve resistance against pathogens in fish [13,16]. However, consumers generally have relatively little awareness of transgenesis and have more negative attitudes toward genetically modified organisms than genome-edited organisms [62], hence the public pushback against transgenic/gene-edited animals is hindering them from reaching the market. Here, we reasonably contend that cathelicidin transgenic catfish would not pose a threat to food safety since: 1) Meat from artificially grown alligators is edible even when consumed raw, and the gut will digest most proteins and inactivate them. 2) Eventually, amino acids rather than proteins are absorbed by humans. 3) Even though the gene sequence is ever-changing in various beings, there are only 20 different types of encoded amino acids that are frequently consumed by humans. In this vein, we are raising attention of potential benefits and risks of our *As-Cath* transgenic catfish by making them transparent to the public.

Nonetheless, scientists and breeders need to be aware of the possible damage that genetically modified fish could cause to the environment and ecosystem [16]. On the one hand, reproductive sterility via genome editing has been attracting the attention of researchers and offering opportunities to reduce environmental risks in aquaculture [62]. On the other hand, representative examples have illustrated that reproductive confinement is promising in model and cultured fish by knocking out/disrupting gonadal development-related genes [23,63-66]. Recently, Qin et al. [19] demonstrated that the reproduction-blocked channel catfish are sterile, and this reproductive confinement can be lifted through hormone therapy with LHRHa. In this study, the dose of 1600 IU/kg HCG coupled with 50 μg/kg LHRHa can restore fecundity at the highest level in comparison to other hormone treatments, but this improvement is not significant from that of 1200 IU/kg HCG. Therefore, a low dose of 1200 IU/kg HCG is recommended for hormone therapy to restore the reproduction of the sterile fish line to reduce costs. In addition to genetically achieving reproductive sterility, well-confined culture systems should be adopted to avoid the escape of mutant/transgenic individuals, especially in the experimental phase of transgenic fish.

## 5. Conclusions

We established a sterile catfish line that confers enhanced resistance to fish pathogens by expressing the cathelicidin protein. Our study has demonstrated that the insertion of the cathelicidin gene at the *LH* locus by harnessing the HA-or ssODN-mediated CRISPR/Cas9 system can be a robust approach to produce sterilized and environmentally-sound fish lines with enhanced disease resistance. Encouragingly, CRISPR/Cas9-mediated KI of AMGs at the reproduction-related loci coupled with hormone therapy could be applied in other commercial fish to increase profits and lower environmental dangers posed by escaped genetic-modified individuals. Notably, even though the desired traits (on-target insertions) can be quickly achieved through CRISPR/Cas9-mediated genome editing, this does not safeguard that we will be able to yield enough non-mosaic P_0_ founders. We contend the genome-editing tool should be used as a complement to existing breeding techniques, not a replacement for them. Hence, a combination of genome editing and conventional selective breeding is required to maximize the benefits of CRISPR/Cas9 tools more effectively in aquatic applications and to hasten the breeding process. In conclusion, this study showed the potential of overexpressing a disease-resistant peptide inserted at a reproduction-related gene using CRISPR/Cas9 in channel catfish, which may provide a strategy of decreasing bacterial disease problems in catfish at the same time reducing environmental risks.

## Acknowledgments

We thank Dr. Eric Peatman for providing the CFX96™ Real-Time System. This project was partially supported by USDA Grant No. G11941 (2018-33522-28769), and Alabama Agricultural Experiment Station grant (AAES-AIR). Jinhai Wang was supported by a scholarship from the China Scholarship Council.

## Appendix A. Supplementary data

Supplementary data to this article can be found from Appendix A and Appendix B.

## References

[1] Doudna JA, Charpentier E. The new frontier of genome engineering with CRISPR-Cas9. Science 2014;346:1258096. https://doi.org/10.1126/science.1258096.

[2] Storici F, Snipe JR, Chan GK, Gordenin DA, Resnick MA. Conservative repair of a chromosomal double-strand break by single-strand DNA through two steps of annealing. Mol Cell Biol 2006;26:7645–57. https://doi.org/10.1128/mcb.00672-06.

[3] Chen F, Pruett-Miller SM, Huang Y, Gjoka M, Duda K, Taunton J, et al. High-frequency genome editing using ssDNA oligonucleotides with zinc-finger nucleases. Nat Methods 2011;8:753–5. https://doi.org/10.1038/nmeth.1653.

[4] Wefers B, Meyer M, Ortiz O, Hrabé de Angelis M, Hansen J, Wurst W, et al. Direct production of mouse disease models by embryo microinjection of TALENs and oligodeoxynucleotides. Proc Natl Acad Sci USA 2013;110:3782–87. https://doi.org/10.1073/pnas.1218721110.

[5] Yoshimi K, Kunihiro Y, Kaneko T, Nagahora H, Voigt B, Mashimo T. ssODN-mediated knock-in with CRISPR-Cas for large genomic regions in zygotes. Nat Commun 2016;7:10431. https://doi.org/10.1038/ncomms10431.

[6] Mehravara M, Shirazia A, Nazaric M, Banand M. Mosaicism in CRISPR/Cas9-mediated genome editing. Dev Biol 2019;445:156–62. https://doi.org/10.1016/j.ydbio.2018.10.008.

[7] Boel A, De Saffel H, Steyaert W, Callewaert B, De Paepe A, Coucke PJ, et al. CRISPR/Cas9-mediated homology-directed repair by ssODNs in zebrafish induces complex mutational patterns resulting from genomic integration of repair-template fragments. Dis Model Mech 2018;11:dmm035352. https://doi.org/10.1242/dmm.035352.

[8] Hisano Y, Sakuma T, Nakade S, Ohga R, Ota S, Okamoto H, et al. Precise in-frame integration of exogenous DNA mediated by CRISPR/Cas9 system in zebrafish. Sci Rep 2015;5:8841. https://doi.org/10.1038/srep08841.

[9] Zhang J-P, Li X-L, Li G-H, Chen W, Arakaki C, Botimer GD, et al. Efficient precise knockin with a double cut HDR donor after CRISPR/Cas9-mediated double-stranded DNA cleavage. Genome Biology 2017;18. https://doi.org/10.1186/s13059-017-1164-8.

[10] Murakami Y, Ansai S, Yonemura A, Kinoshita M. An efficient system for homology-dependent targeted gene integration in medaka (Oryzias latipes). Zoological Lett 2017;3:10. https://doi.org/10.1186/s40851-017-0071-x.

[11] Ledford H. Salmon approval heralds rethink of transgenic animals. Nature 2015;527:417–18. https://doi.org/10.1038/527417a.

[12] Waltz E. First genetically engineered salmon sold in Canada. Nature 2017;548:148–48. https://doi.org/10.1038/nature.2017.22116.

[13] Wang J, Su B, Dunham RA. Genome-wide identification of catfish antimicrobial peptides: A new perspective to enhance fish disease resistance. Rev Aquac 2022a;14:2002–22. https://doi.org/10.1111/raq.12684.

[14] Xing D, Su B, Bangs M, Li S, Wang J, Bern L, et al. CRISPR/Cas9-mediate knock-in method can improve the expression and effect of transgene in P1 generation of channel catfish (Ictalurus punctatus). Aquaculture. 2022a;560:738531. https://doi.org/10.1016/j.aquaculture.2022.738531.

[15] Xing D, Su B, Li S, Bangs M, Creamer D, Coogan M, et al. CRISPR/Cas9-mediated transgenesis of the Masu salmon (Oncorhynchus masou) elovl2 gene improves n-3 fatty acid content in channel catfish (Ictalurus punctatus). Mar Biotechnol 2022b;24:513–23. https://doi.org/10.1007/s10126-022-10110-6.

[16] Dunham RA, Su B. Genetically Engineered Fish: Potential Impacts on Aquaculture, Biodiversity, and the Environment. In: Chaurasia A, Hawksworth DL, Pessoa de Miranda M, editors. GMOs: Implications for Biodiversity Conservation and Ecological Processes, Cham: Springer International Publishing; 2020, p. 241–75. https://doi.org/10.1007/978-3-030-53183-6_11.

[17] Blix TB, Dalmo RA, Wargelius A, Myhr AI. Genome editing on finfish: Current status and implications for sustainability. Rev Aquac 2021;13:2344–63. https://doi.org/10.1111/raq.12571.

[18] Yang Z, Yu Y, Tay YX, Yue GH. Genome editing and its applications in genetic improvement in aquaculture. Rev Aquac 2022;14:178–91. https://doi.org/10.1111/raq.12591.

[19] Qin G, Qin Z, Lu C, Ye Z, Elaswad A, Bangs M, et al. Gene editing of the catfish gonadotropin-releasing hormone gene and hormone therapy to control the reproduction in channel catfish, Ictalurus punctatus. Biology (Basel) 2022;11:649. https://doi.org/10.3390/biology11050649.

[20] Grier HJ. Cellular organization of the testis and spermatogenesis in fishes. Am Zool 1981;21:345–57. https://doi.org/10.1093/icb/21.2.345.

[21] Yamaguchi Y, Nagata J, Nishimiya O, Kawasaki T, Hiramatsu N, Todo T. Molecular characterization of fshb and lhb subunits and their expression profiles in captive white-edged rockfish, Sebastes taczanowskii. Comp Biochem Physiol A: Mol Integr Physiol 2021;261:111055. https://doi.org/10.1016/j.cbpa.2021.111055.

[22] Chu L, Li J, Liu Y, Hu W, Cheng CHK. Targeted gene disruption in zebrafish reveals noncanonical functions of LH signaling in reproduction. Mol Endocrinol 2014;28:1785–95. https://doi.org/10.1210/me.2014-1061.

[23] Qin Z, Li Y, Su B, Cheng Q, Ye Z, Perera DA, et al. Editing of the luteinizing hormone gene to sterilize channel catfish, Ictalurus punctatus, using a modified zinc finger nuclease technology with electroporation. Mar Biotechnol 2016;18:255–63. https://doi.org/10.1007/s10126-016-9687-7.

[24] Akira S, Uematsu S, Takeuchi O. Pathogen recognition and innate immunity. Cell 2006;124:783–801. https://doi.org/10.1016/j.cell.2006.02.015.

[25] Wang G, Li X, Wang Z. APD3: the antimicrobial peptide database as a tool for research and education. Nucleic Acids Res 2016;44:D1087–93. https://doi.org/10.1093/nar/gkv1278.

[26] Wang J, Wilson AE, Su B, Dunham RA. Functionality of dietary antimicrobial peptides in aquatic animal health: Multiple meta-analyses. Anim Nutr 2022b. https://doi.org/10.1016/j.aninu.2022.10.001.

[27] Mookherjee N, Anderson MA, Haagsman HP, Davidson DJ. Antimicrobial host defence peptides: functions and clinical potential. Nat Rev Drug Discov 2020;19:311–32. https://doi.org/10.1038/s41573-019-0058-8.

[28] Hilchie AL, Wuerth K, Hancock REW. Immune modulation by multifaceted cationic host defense (antimicrobial) peptides. Nat Chem Biol 2013;9:761–68. https://doi.org/10.1038/nchembio.1393.

[29] Chen Y, Cai S, Qiao X, Wu M, Guo Z, Wang R, et al. As-CATH1-6, novel cathelicidins with potent antimicrobial and immunomodulatory properties from Alligator sinensis, play pivotal roles in host antimicrobial immune responses. Biochem J 2017;474:2861–85. https://doi.org/10.1042/BCJ20170334.

[30] Simora RMC, Li S, Abass NY, Terhune JS, Dunham RA. Cathelicidins enhance protection of channel catfish, Ictalurus punctatus, and channel catfish ♀ × blue catfish, Ictalurus furcatus ♂ hybrid catfish against Edwarsiella ictaluri infection. J Fish Dis 2020;43:1553–62. https://doi.org/10.1111/jfd.13257.

[31] Simora RMC, Wang W, Coogan M, El Husseini N, Terhune JS, Dunham RA. Effectiveness of cathelicidin antimicrobial peptide against Ictalurid catfish bacterial pathogens. J Aquat Anim Health 2021;33:178–89. https://doi.org/10.1002/aah.10131.

[32] Liu Z, Liu S, Yao J, Bao L, Zhang J, Li Y, et al. The channel catfish genome sequence provides insights into the evolution of scale formation in teleosts. Nat Commun 2016;7:11757. https://doi.org/10.1038/ncomms11757.

[33] Mosimann C, Kaufman CK, Li P, Pugach EK, Tamplin OJ, Zon LI. Ubiquitous transgene expression and Cre-based recombination driven by the ubiquitin promoter in zebrafish. Development 2011;138:169–77. https://doi.org/10.1242/dev.059345.

[34] Bae S, Park J, Kim J-S. Cas-OFFinder: a fast and versatile algorithm that searches for potential off-target sites of Cas9 RNA-guided endonucleases. Bioinformatics 2014;30:1473–75. https://doi.org/10.1093/bioinformatics/btu048.

[35] Elaswad A, Khalil K, Ye Z, Liu Z, Liu S, Peatman E, et al. Effects of CRISPR/Cas9 dosage on TICAM1 and RBL gene mutation rate, embryonic development, hatchability and fry survival in channel catfish. Sci Rep 2018;8:16499. https://doi.org/10.1038/s41598-018-34738-4.

[36] Khalil K, Elayat M, Khalifa E, Daghash S, Elaswad A, Miller M, et al. Generation of myostatin gene-edited channel catfish (Ictalurus punctatus) via zygote injection of CRISPR/Cas9 system. Sci Rep 2017;7:7301. https://doi.org/10.1038/s41598-017-07223-7.

[37] Armstrong JB, Malacinski GM. Developmental Biology of the Axolotl. New York: Oxford University Press; 1989.

[38] Qiu P, Shandilya H, D’Alessio JM, O’Connor K, Durocher J, Gerard GF. Mutation detection using Surveyor™ nuclease. Biotechniques 2004;36:702–7. https://doi.org/10.2144/04364PF01.

[39] Coogan M, Alston V, Su B, Khalil K, Elaswad A, Khan M, et al. CRISPR/Cas-9 induced knockout of myostatin gene improves growth and disease resistance in channel catfish (Ictalurus punctatus). Aquaculture 2022;557:738290. https://doi.org/10.1016/j.aquaculture.2022.738290.

[40] Davis KB. Age at puberty of channel catfish, Ictalurus punctatus, controlled by thermoperiod. Aquaculture 2009;292:244–47. https://doi.org/10.1016/j.aquaculture.2009.04.023.

[41] Menchaca A, Dos Santos-Neto PC, Souza-Neves M, Cuadro F, Mulet AP, Tesson L, et al. Otoferlin gene editing in sheep via CRISPR-assisted ssODN-mediated homology directed repair. Sci Rep 2020;10:5995. https://doi.org/10.1038/s41598-020-62879-y.

[42] Kan Y, Ruis B, Takasugi T, Hendrickson EA. Mechanisms of precise genome editing using oligonucleotide donors. Genome Res 2017;27:1099–111. https://doi.org/10.1101/gr.214775.116.

[43] Byrne SM, Ortiz L, Mali P, Aach J, Church GM. Multi-kilobase homozygous targeted gene replacement in human induced pluripotent stem cells. Nucleic Acids Res 2015;43:e21. https://doi.org/10.1093/nar/gku1246.

[44] Simora RMC, Xing D, Bangs MR, Wang W, Ma X, Su B, et al. CRISPR/Cas9[mediated knock[in of alligator cathelicidin gene in a non[coding region of channel catfish genome. Sci Rep 2020;10:22271. https://doi.org/10.1038/s41598-020-79409-5.

[45] McGaw C, Garrity AJ, Munoz GZ, Haswell JR, Sengupta S, Keston-Smith E, et al. Engineered Cas12i2 is a versatile high-efficiency platform for therapeutic genome editing. Nat Commun 2022;13:2833 https://doi.org/10.1038/s41467-022-30465-7.

[46] Kelly JJ, Saee-Marand M, Nyström NN, Evans MM, Chen Y, Martinez FM, et al. Safe harbor-targeted CRISPR-Cas9 homology-independent targeted integration for multimodality reporter gene-based cell tracking. Sci Adv 2021;7: eabc3791. https://doi.org/10.1126/sciadv.abc3791

[47] Wang C, Qu Y, Cheng JKW, Hughes NW, Zhang Q, Wang M, et al. dCas9-based gene editing for cleavage-free genomic knock-in of long sequences. Nat Cell Biol 2022;24:268–78. https://doi.org/10.1038/s41556-021-00836-1.

[48] Ran FA, Hsu PD, Wright J, Agarwala V, Scott DA, Zhang F. Genome engineering using the CRISPR-Cas9 system. Nat Protoc 2013;8:2281–308. https://doi.org/10.1038/nprot.2013.143.

[49] Heigwer F, Kerr G, Boutros M. E-CRISP: fast CRISPR target site identification. Nat Methods 2014;11:122–23. https://doi.org/10.1038/nmeth.2812.

[50] Zhang X-H, Tee LY, Wang X-G, Huang Q-S, Yang S-H. Off-target effects in CRISPR/Cas9-mediated genome engineering. Mol Ther Nucleic Acids 2015;17:e264. https://doi.org/10.1038/mtna.2015.37.

[51] Pattanayak V, Lin S, Guilinger JP, Ma E, Doudna JA, Liu DR. High-throughput profiling of off-target DNA cleavage reveals RNA-programmed Cas9 nuclease specificity. Nat Biotechnol 2013;31:839–43. https://doi.org/10.1038/nbt.2673.

[52] Cho SW, Kim S, Kim Y, Kweon J, Kim HS, Bae S, et al. Analysis of off-target effects of CRISPR/Cas-derived RNA-guided endonucleases and nickases. Genome Res 2014;24:132–41. https://doi.org/10.1101/gr.162339.113.

[53] Shen B, Zhang J, Wu H, Wang J, Ma K, Li Z, et al. Generation of gene-modified mice via Cas9/RNA-mediated gene targeting. Cell Res 2013;23:720–23. https://doi.org/10.1038/cr.2013.46

[54] Iyer V, Shen B, Zhang W, Hodgkins A, Keane T, Huang X, et al. Off-target mutations are rare in Cas9-modified mice. Nat Methods 2015;12:479. https://doi.org/10.1038/nmeth.3408.

[55] Wang T, Wei JJ, Sabatini DM, Lander, ES. Genetic screens in human cells using the CRISPR-Cas9 system. Science 2014;343:80–84. https://doi.org/10.1126/science.1246981.

[56] Li D, Qiu Z, Shao Y, Chen Y, Guan Y, Liu M, et al. Heritable gene targeting in the mouse and rat using a CRISPR-Cas system. Nat Biotechnol 2013;31:681–83. https://doi.org/10.1038/nbt.2661.

[57] Li W, Teng F, Li T, Zhou Q. Simultaneous generation and germline transmission of multiple gene mutations in rat using CRISPR-Cas systems. Nat Biotechnol 2013;31:684. https://doi.org/10.1038/nbt.2652.

[58] Oliver D, Yuan S, McSwiggin H, Yan W. Pervasive genotypic mosaicism in founder mice derived from genome editing through pronuclear injection. PLOS One 2015;10: e0129457. https://doi.org/10.1371/journal.pone.0129457.

[59] Raveux A, Vandormael-Pournin S, Cohen-Tannoudji M. Optimization of the production of knock-in alleles by CRISPR/Cas9 microinjection into the mouse zygote. Sci Rep 2017;7:42661. https://doi.org/10.1038/srep42661.

[60] Jao L-E, Wente SR, Chen W. Efficient multiplex biallelic zebrafish genome editing using a CRISPR nuclease system. Proc Natl Acad Sci USA 2013;110:13904–909. https://doi.org/10.1073/pnas.1308335110.

[61] Auer TO, Duroure K, De Cian A, Concordet JP, Del Bene F. Highly efficient CRISPR/Cas9-mediated knock-in in zebrafish by homology-independent DNA repair. Genome Res 2014;24:142–53. https://doi.org/10.1101/gr.161638.113.

[62] Hallerman EM, Dunham R, Houston RD, Walton M, Wargelius A, Wray-Cahen D. Towards production of genome-edited aquaculture species. Rev Aquac 2022;1–5. https://doi.org/10.1111/raq.12739.

[63] Wargelius A, Leininger S, Skaftnesmo KO, Kleppe L, Andersson E, Taranger GL, et al. Dnd knockout ablates germ cells and demonstrates germ cell independent sex differentiation in Atlantic salmon. Sci Rep. 2016;6:21284. https://doi.org/10.1038/srep21284.

[64] Gay S, Bugeon J, Bouchareb A, Henry L, Delahaye C, Legeai F, et al. MiR-202 controls female fecundity by regulating medaka oogenesis. PLoS Genet 2018;14:e1007593. https://doi.org/10.1371/journal.pgen.1007593.

[65] Su B, Peatman E, Shang M, Thresher R, Grewe P, Patil JG, et al. Expression and knockdown of primordial germ cell genes, vasa, nanos and dead end in common carp (Cyprinus carpio) embryos for transgenic sterilization and reduced sexual maturity. Aquaculture 2014;S72–S84:420–21. https://doi.org/10.1016/j.aquaculture.2013.07.008.

[66] Su B, Shang M, Grewe PM, Patil JG, Peatman E, Perera DA, et al. Suppression and restoration of primordial germ cell marker gene expression in channel catfish, Ictalurus punctatus, using knockdown constructs regulated by copper transport protein gene promoters: Potential for reversible transgenic sterilization. Theriogenology 2015;84:1499–512. https://doi.org/10.1016/j.theriogenology.2015.07.037.

